# QTL analysis reveals an oligogenic architecture of a rapidly adapting trait during the European invasion of common ragweed

**DOI:** 10.1101/2022.02.24.481758

**Authors:** Diana Prapas, Romain Scalone, Jacqueline Lee, Kristin A Nurkowski, Sarah Bou-assi, Loren Rieseberg, Paul Battlay, Kathryn A Hodgins

## Abstract

Biological invasions offer a unique opportunity to investigate evolution over contemporary time-scales. Rapid adaptation to local climates during range expansion can be a major determinant of invasion success, yet fundamental questions remain about its genetic basis. This study sought to investigate the genetic basis of climate adaptation in invasive common ragweed (*Ambrosia artemisiifolia*). Flowering time adaptation is key to this annual species’ invasion success, so much so that it has evolved repeated latitudinal clines in size and phenology across its native and introduced ranges despite high gene flow among populations. Here, we produced a high-density linkage map (4,493 SNPs) and paired this with phenotypic data from an F2 mapping population (n=336) to identify one major and two minor quantitative trait loci (QTL) underlying flowering time and height differentiation in this species. Within each QTL interval, several candidate flowering time genes were also identified. Notably, the major flowering time QTL detected in this study was found to overlap with a previously identified haploblock (putative inversion). Multiple genetic maps of this region identified evidence of suppressed recombination in specific genotypes, consistent with inversions. These discoveries support the expectation that a concentrated genetic architecture with fewer, larger and more tightly-linked alleles should underlie rapid local adaptation during invasion, particularly when divergently-adapting populations experience high-levels of gene flow.

## Introduction

While invasive species often have disastrous economic or ecological impacts, and thereby are important subjects for applied research, they remain a fascinating evolutionary phenomenon. Exotic and weedy species are highly successful colonisers that can rapidly spread through diverse habitats, responding to changing environmental conditions and novel selection pressures along the way (Bock et al., 2015; Hodgins, Bock, & Rieseberg, 2018). How they are able to thrive in foreign environments is of great interest, especially considering the reductions in genetic diversity that can often accompany long-distance range expansion (Franks & Munshi-South, 2014; Sax & Brown, 2000). These features make biological invasions a natural experiment that can be used to investigate adaptation over contemporary time-scales (Dlugosch, Anderson, Braasch, Cang, & Gillette, 2015; Gilchrist & Lee, 2007; McGoey, Hodgins, & Stinchcombe, 2020; Sakai et al., 2001; Westley, 2011). Applying an evolutionary lens to invasion biology is also critical to understanding invasion trajectories and to what extent they might be predictable in light of anthropogenic land use and climate change (van Boheemen & Hodgins, 2020). Hence, what is notoriously viewed as an economic and ecological problem also provides a promising evolutionary research opportunity (Callaway & Maron, 2006).

A large body of evolutionary research points to local adaptation as being an important mechanism for invasion success (Colautti & Barrett, 2013; Sherrard & Maherali, 2012). When an invasive species colonises new regions, the invaders must contend with environmental variation in order to establish (Colautti & Lau, 2015). It is this environmental heterogeneity that can impose selection to favour traits differentially and push populations towards divergent trait optima, resulting in local adaptation (Ågren, Oakley, Lundemo, & Schemske, 2017; Hereford, 2009; Hoban et al., 2016; Orr, 2005). For local adaptation to occur however, gene flow should not overwhelm the effects of selection and genotypes conferring high fitness in one environment must bear a cost in a contrasting environment (Savolainen, Lascoux, & Merilä, 2013). This distinct signature of local adaptation has been detected during numerous reciprocal transplant experiments, where local genotypes have a home-site advantage over nonlocal genotypes (Hoban et al., 2016; Kawecki & Ebert, 2004). Many studies have also demonstrated the prevalence of local adaptation in a range of taxa, including animals (Fraser, Weir, Bernatchez, Hansen, & Taylor, 2011; Hereford, 2009) and plants (Leimu & Fischer, 2008). However in invasive species, there is extensive evidence of rapid local adaptation (Colautti & Barrett, 2013; Franks, Wheeler, & Goodnight, 2012; Oduor, Leimu, & van Kleunen, 2016; Sultan, Horgan-Kobelski, Nichols, Riggs, & Waples, 2013), revealing that evolutionary changes can take place on ecological timescales (Colautti & Lau, 2015), much more rapidly than traditionally thought.

Clinal variation in fitness-related traits is a striking manifestation of local adaptation (Savolainen et al., 2013). These patterns of population differentiation are shaped by climatic factors that are associated with latitudinal or altitudinal gradients (Bruelheide & Heinemeyer, 2002; Chun, Le Corre, & Bretagnolle, 2011). Whilst trait clines can occur non-adaptively via neutral evolutionary processes (Colautti & Lau, 2015), a growing number of empirical studies have shown that adaptive responses to heterogeneous climatic pressures often explain the emergence of geographic clines *in situ* (Keller & Taylor, 2008; Santangelo, Johnson, & Ness, 2018). Oftentimes, these clines are the result of evolutionary trade-offs (Colautti & Barrett, 2010). For instance, a trade-off between growth and reproduction is frequently reported to shape adaptive trait divergence in many plant populations (Colautti & Barrett, 2010; Griffith & Watson, 2005; Grime, 1977). This trade-off directly relates to reproductive initiation or flowering time (a critical life-history trait and a key component of fitness in flowering plants), where the best strategy depends on local environmental conditions (Anderson, Willis, & Mitchell-Olds, 2011; Yan, Wang, Chan, & Mitchell-Olds, 2021). In temperate environments, at the cost of reduced size, plants in short growing seasons tend to flower earlier to ensure successful reproduction before the onset of harsh fall frosts (Anderson et al., 2011; Grime, 1977; Stinson, Wheeler, Record, & Jennings, 2018). Contrastingly, where the growing season is longer, plants can afford to delay their flowering in exchange for enhanced growth, fecundity and competitiveness (Colautti & Barrett, 2010; Kralemann, Scalone, Andersson, & Hennig, 2018; Stinson et al., 2018). This latitudinal cline in plant size and flowering time has been documented in several invasive plant species, such as *Lythrum salicaria* (Colautti & Barrett, 2013), *Medicago polymorpha* (Helliwell et al., 2018), and *Ambrosia artemisiifolia* (Hodgins & Rieseberg, 2011; McGoey et al., 2020; Scalone et al., 2016; van Boheemen, Atwater, & Hodgins, 2019).

Despite these preliminary insights, the genetic basis of climate adaptation remains poorly understood. Primarily, there is a large knowledge gap concerning the genetic architecture underlying local adaptation during invasion. In most cases, ecologically relevant traits that confer local adaptation are quantitative; genetic variants responsible for such quantitative trait variation are located in genomic regions known as quantitative trait loci (QTL) (Collard, Jahufer, Brouwer, & Pang, 2005). However, despite the perceived importance of quantitative traits in local adaptation, research has been more focussed on characterizing discrete traits with simple Mendelian modes of inheritance (Savolainen et al., 2013). Therefore, our understanding of invasion dynamics from a quantitative genetics perspective is limited. Specifically, the number, the distribution and the effect-sizes of QTL that contribute to adaptive trait variation are unclear, especially in the invasion context. While Fisher’s infinitesimal model suggests that quantitative trait variance should be controlled by many loci of small-effect (Fisher, 1919; Louthan & Kay, 2011), a genetic architecture with contributions of loci of larger phenotypic effect is expected to benefit a population that is initially far from its phenotypic optima (MacPherson & Nuismer, 2017; Orr, 1998), which is typical of an invader undergoing rapid local adaptation (Bock et al., 2015). During invasion, theory predicts that a combination of both large and small-effect loci will more frequently contribute to evolutionary rescue by working synergistically to maximise fitness (Gomulkiewicz, Holt, Barfield, & Nuismer, 2010). Large-effect loci will also preferentially contribute to adaptive divergence when migration is high and/or drift is strong (Hodgins & Yeaman, 2019; Yeaman & Whitlock, 2011). Additionally, clusters of small-effect loci can effectively act as a large-effect QTL if recombination between them is sufficiently low (Yeaman & Whitlock, 2011), such as in the case of a chromosomal inversion. Both gene flow and drift might be important in promoting more concentrated genetic architectures (i.e., fewer loci of large effect, or multiple linked loci) in wide-ranging invasive species. In their native ranges, species connected by high gene flow over heterogeneous environments may evolve concentrated architectures over the long term. Upon introduction, strong drift may filter out weakly-selected loci during invasion bottlenecks (Dlugosch et al., 2015), and distant phenotypic optima will promote the contribution of large-effect loci to adaptive trait divergence. More empirical evidence is, however, needed to help evaluate these theories.

Common ragweed (*Ambrosia artemisiifolia*) is an annual weed native to North America that has successfully invaded a range of environments across the globe (Chauvel, Dessaint, Cardinal-Legrand, & Bretagnolle, 2006; Essl et al., 2015; Smith, Cecchi, Skjøth, Karrer, & Šikoparija, 2013; van Boheemen & Hodgins, 2020), and is an important study system for investigating the genetic basis of climate adaptation. Firstly, due to its allelopathic effects on several major crop species and its highly allergenic pollen, common ragweed is of major agricultural and human-health concern (Bassett & Crompton, 1975; Laaidi, Laaidi, Besancenot, & Thibaudon, 2003; Tokarska-Guzik et al., 2011). Contemporary climate change is expected to extend ragweed’s pollen season and accelerate its range expansion at the northern margins of its current distribution (Chapman, Scalone, Štefanić, & Bullock, 2017; Cunze, Leiblein, & Tackenberg, 2013; Ziska et al., 2011). Understanding how it may evolve and spread in the future is crucial to better managing the incidence of hay-fever and preventing further yield losses. Secondly, the species has evolved parallel latitudinal clines in size and phenology in its native North American range and its introduced ranges of Europe (Chun et al., 2011), Asia (Li, She, Zhang, & Liao, 2015) and Australia (van Boheemen et al., 2019, 2017). Remarkably, the clines observed in the introduced ranges evolved in a mere 100-150 years, signifying rapid adaptation in this species (Hodgins & Rieseberg, 2011; Leiblein-Wild & Tackenberg, 2014; Scalone et al., 2016; van Boheemen et al., 2019). There is further evidence to suggest that this clinal variation in flowering time is the result of adaptive rather than neutral processes (McGoey et al., 2020; van Boheemen et al., 2019). Hence, this species offers an important opportunity to study the genetic basis of rapid, local adaptation during range expansion.

The main objective of this study was to elucidate the genetic basis of climate adaptation in common ragweed. Specifically, we aimed to identify the underlying genetic architecture of flowering time and plant height, key adaptive traits in this annual species, by conducting QTL mapping. We first developed a linkage map using F2 progeny derived from two experimental crosses between an early flowering, introduced-range parent, and a late flowering, native-range parent. All plants were then genotyped using Genotype-by-Sequencing (GBS), and phenotyped under controlled environmental conditions. Based on the evolutionary theory discussed above, we expected to identify an oligogenic trait architecture, with some larger-effect loci contributing to flowering time divergence. This is because common ragweed is an outcrossing wind-pollinated species with high population connectivity, yet strong flowering time differentiation, even in recently invaded ranges, consistent with rapid and recent local adaptation (Chun et al., 2011; McGoey et al., 2020; van Boheemen et al., 2019). We also anticipated that the flowering time QTL would colocalize with height QTL, due to the expected genetic correlation between flowering time and height in this species.

## Methods

### Study species

Common ragweed (*Ambrosia artemisiifolia*) is a monoecious annual in the Asteraceae family (Bassett & Crompton, 1975; Smith et al., 2013). This outcrossing plant is a globally invasive species that has aggressively spread from its native North American range to many regions of the world (Friedman & Barrett, 2008). This species thrives in disturbed habitats and is primarily found growing alongside roads and in cultivated areas where competition from other plants is limited (Bassett & Crompton, 1975). It is widely recognised as a noxious agricultural weed due to its allelopathic effects on crop species such as soybean and maize (Weaver, 2001). Despite low genetic structure and high gene flow across much of the species’ range (McGoey & Stinchcombe, 2021; van Boheemen et al., 2017), the species has locally adapted and evolved parallel latitudinal clines in size and phenology where high-latitudinal populations flower early when small and low-latitudinal populations flower later when tall (Chun et al., 2011; Hodgins & Rieseberg, 2011; Li et al., 2015; van Boheemen et al., 2019). Its wind-spread pollen can travel several thousands of kilometres and can induce allergic rhinitis in human populations, costing millions of dollars in medical treatment each year (Taramarcaz, Lambelet, Clot, Keimer, & Hauser, 2005).

### Experimental crosses

We generated experimental mapping populations where early-flowering individuals from the northern part of the introduced European range (Drebkau, Germany N51°38’21’’ E14°11’50’’) were crossed with late-flowering individuals from the central part of the native North American range (Lexington, KY, USA N38°01’ W84°33’) to produce F1 offspring. *Ambrosia artemisiifolia* seeds of both populations were stratified on moist filter paper for 4 weeks and subsequently placed in a growth chamber providing 16-hour light of 50 μmol m^-2^ s^-1^ at 27 °C and 8-hour darkness at 15 °C to induce germination. After germination, seedlings were transplanted to pots with pumice (0.5 mm<diameter>2.8 mm; Hekla Green, Bara Mineraler, Bara, Sweden), and moved into a climate chamber with short-day conditions (12-hour light of 280μmol m^-2^ s^-1^ at 25°C; 12-hour darkness at 15°C; humidity at 70%). Plants were watered with nutrient-enriched water (2 ml l–1 of Wallco växtnäring 53-10-43+micro from Cederroth, Upplands Väsby, Sweden), if necessary. The position of the plants in the growth chamber was randomized daily. The induction of the germination was executed at different moments for each seed population (the seeds of the late-flowering population were induced before the seeds of the early-flowering population, according to the flowering time differences observed in (Scalone et al., 2016)), in order to “synchronise” them. When two plants - one of each population - were presenting similar growth and flowering development, they were both placed in the same individual cabinet, parameterized with identical temperature and light conditions and cycles than the growth chamber. The use of an individual and closed cabinet (located in different sectors of the Uppsala BioCenter) prevented pollen contamination among crosses. During the flowering period of each crossing pair, the maternal plant was emasculated to prevent self pollination (although the species is self-incompatible) (Friedman & Barrett, 2008). For each cross/cabinet, two pollination methods were used daily: first, the yellowish pollen of the paternal plant was collected with an individual brush to pollinate female flowers of the maternal plant. Then, secondarily, before closing the 2-door system of the cabinet and after the “brush”-pollination session, the paternal plant was gently slapped in order to generate/spread a “pollen cloud” inside the hermetic cabinet. At the end of the flowering period, F1 seeds were collected and stratified. Then, similar experimental conditions and techniques were used to produce F2 populations from these F1 seeds. Two F1 siblings from each set of parents were then inter-crossed to produce segregating F2 populations. F2 plants were grown in a growth chamber with a 12-hour daylight/darkness cycle where leaves material of each F2-plants was collected for DNA extraction and the following phenotypes were scored: Initial budding time (number of days following germination at which a pale green budding point appeared), 1cm budding time (number of days following germination at which the bud was greater than 1cm), male and female flowering times (days following germination at which anthers/pistils first appeared) and plant height (including stem and terminal male inflorescence). Three experimental mapping families (named pink, yellow and orange) were created from six different grand-parental plants (three obtained from the early flowering German population and three obtained from the late flowering Kentucky population). Leaf material of grand- and parental plants were collected for DNA extraction.

We generated a fourth mapping population at the University of British Columbia. This F1 mapping population consisted of a cross between a single individual from North America (AA19_3_7, ND, USA N46°17’ W103°55’), which was used in the creation of a reference genome assembly (Bieker et al., 2022), and a single individual from France (FR8-26-8, France N44°12’ E4°15’). The two plants were grown alone in a growth chamber to facilitate crossing between the two wind pollinated individuals and prevent pollen contamination from other individuals, and the maternal plant was emasculated to prevent self pollination (although the plants are self-incompatible) (Friedman & Barrett, 2008). We collected the seeds from FR8-26-8. Following this, we grew offspring from the cross in a glasshouse at Monash University and their leaf tissue was harvested for DNA extraction. Tissue was preserved from the parents and offspring by drying using silica gel.

### Sequencing and data processing

We obtained marker loci using DNA extraction and genotype-by-sequencing (GBS) protocols described in van Boheemen *et al.* (2019). The libraries were sequenced on a HiSeq2500 (125 bp PE) at the Génome Québec Innovation Centre on four lanes resulting in 625 million paired end reads. Reads were demultiplexed with Stacks process_radtags (Catchen, Hohenlohe, Bassham, Amores, & Cresko, 2013). We trimmed and filtered reads using Fastx (http://hannonlab.cshl.edu/fastx_toolkit), allowing for Q-score of 20 or higher for ≥90% of the reads. We then aligned the FASTA files to a new chromosome-level reference genome (Battlay *et al.* in prep.) using BWA-mem (Heng Li & Durbin, 2010). SAMtools (Heng Li et al., 2009) was used to call SNPs on each scaffold and the resulting files were concatenated to produce a single VCF file retaining only SNPs genotyped with a minimum base quality score of 10. This file was further filtered with VCFtools (Danecek et al., 2011) to remove indels and exclude sites on the basis of the proportion of missing data (tolerating 25% missing data), leaving only biallelic SNPs with a mean depth of 10 and minimum quality threshold score of ≥20. SNPs that had minor allele frequencies <5% were also discarded. The genotype data from two mapping families (the yellow and pink families) were used for linkage map construction for the QTL analysis, as the third family did not produce a sufficient sample size of genotyped plants. The F1 mapping population was not phenotyped and not used in the construction of the linkage map for the QTL analysis.

### Linkage map construction

Using SNP data from two mapping families (pink and yellow families - four grandparents, four F1-parents and 444 F2 offspring), we constructed an integrated linkage map using the software package Lep-MAP3 (Rastas, 2017). Firstly, the ParentCall2 module identified 26,921 informative markers based on pedigree information and imputed parental genotypes. The Filtering2 module was then used to remove markers that exhibited high segregation distortion (*dataTolerance=0.001*). Next, markers were assigned into linkage groups (LGs) using the SeparateChromosomes2 module which computes all pairwise LOD scores between markers and groups those with a LOD score higher than a user given parameter. It is important to note that the grouping of two families can be inherently difficult as each marker can be informative only on one family (Rastas, 2017). Hence, markers were grouped separately using markers from the pink family first via parameter *families=Pink*. Clustering appeared to be optimal with a LOD score of 15 (*lodLimit=15*) and when the minimum number of markers per LG was set to 25 (*sizeLimit=25*). Specifically, these parameters produced 18 major linkage groups, showing good correspondence with common ragweed’s karyotype (2n=36; Essl *et al*. 2015).

The data containing the common markers for both yellow and pink families were then used to assign the remaining singular markers to the existing linkage groups using JoinSingles2All (*lodLimit=14*). The markers on LG1 were further split to even out the size distribution of linkage groups using SeparateChromosomes2 (*lg=1, lodLimit=20* and *sizeLimit=25*) and the JoinSingles2All map file. Markers within each LG were then ordered using the OrderMarkers2 module (*outputPhasedData=1*, *sexAveraged=1*). As the physical positions of markers from the reference genome were known, this information was used to correct marker order via parameters *evaluateOrder=order.txt* and *improveOrder=0, sexAveraged=1* where order.txt contained the markers in their physical order. Additionally, markers that inflated the length of LGs (>25 cM) or caused LGs to be spread across multiple scaffolds were removed. These orders were evaluated again using an additional parameter (*grandparentPhase=1*) to code genotypes according to grandparental inheritance. To obtain a maximum number of markers, the two evaluated map outputs were matched using phasematch.awk script provided with Lep-MAP3. Finally, the map2genotypes.awk script of Lep-MAP3 was used to convert these phased outputs to fully informative genotype data.

As genomic analysis suggested that some of the identified QTL intervals fell within two putative inversions (haploblocks) on Scaffold 27 (Battlay *et al*. in prep.), we developed sexspecific genetic maps for this scaffold and in each mapping population (pink, yellow and F1 mapping population) to determine if recombination rates were suppressed within these regions in some genotypes. Genotype specific suppression of recombination in the regions would therefore be consistent with inversions rather than general reductions of recombination in the region. Linkage map construction was carried out using the physical order of the markers from Scaffold 27. Genetic distance (cM) was plotted against physical position along the chromosome for each map and the intervals of the QTL and the boundaries of the haploblocks were visualised and inspected for reduced recombination compared to the rest of the scaffold.

### QTL analysis

QTL analysis was performed in R (version 4.0.5) using the R/qtl package (Broman, Wu, Sen, & Churchill, 2003). Since a minimum of 200 individuals are needed to harness sufficient phenotypic power (Beavis, 2019; Li, Hearne, Bänziger, Li, & Wang, 2010), we only conducted QTL scans on the yellow family as it had a larger sample size (n=336) compared to the pink family (n=108). To input the data, yellow F2s were treated as an outbreeding full-sibling population and were therefore read-in as a four-way cross.

### Phenotype and genotype data checks

QTL analysis relies on the assumption that phenotypes follow a normal distribution (Broman & Sen, 2009). To check for adherence to normality, histograms of each phenotype were plotted accordingly. Height was normally distributed, however the remaining budding and flowering time traits were slightly skewed (Figure 1) and so were log-transformed. Prior to QTL analysis, further linkage map and genotype diagnostic tests were conducted using R/qtl. Firstly, markers were checked again for segregation distortion (a departure from 1:1:1:1 in a 4-way cross) using a χ^2^ test. A total of 398 markers on LGs 2, 3, 4, 7, 8, 13 and 18 showed extreme segregation distortion and so these markers were removed. To test for unusually similar genotypes (indicating potential sample mix-ups), the proportion of identical genotypes for each pair of individuals were calculated. These were all below 80%, suggesting no unusually similar pairs. Next, pairwise recombination fractions between all pairs of markers were estimated to check marker assignment to LGs. The obtained recombination heat map showed that some markers on LG14 and LG15 appeared to be linked and so these markers (~100) were omitted.

**Figure 1.**
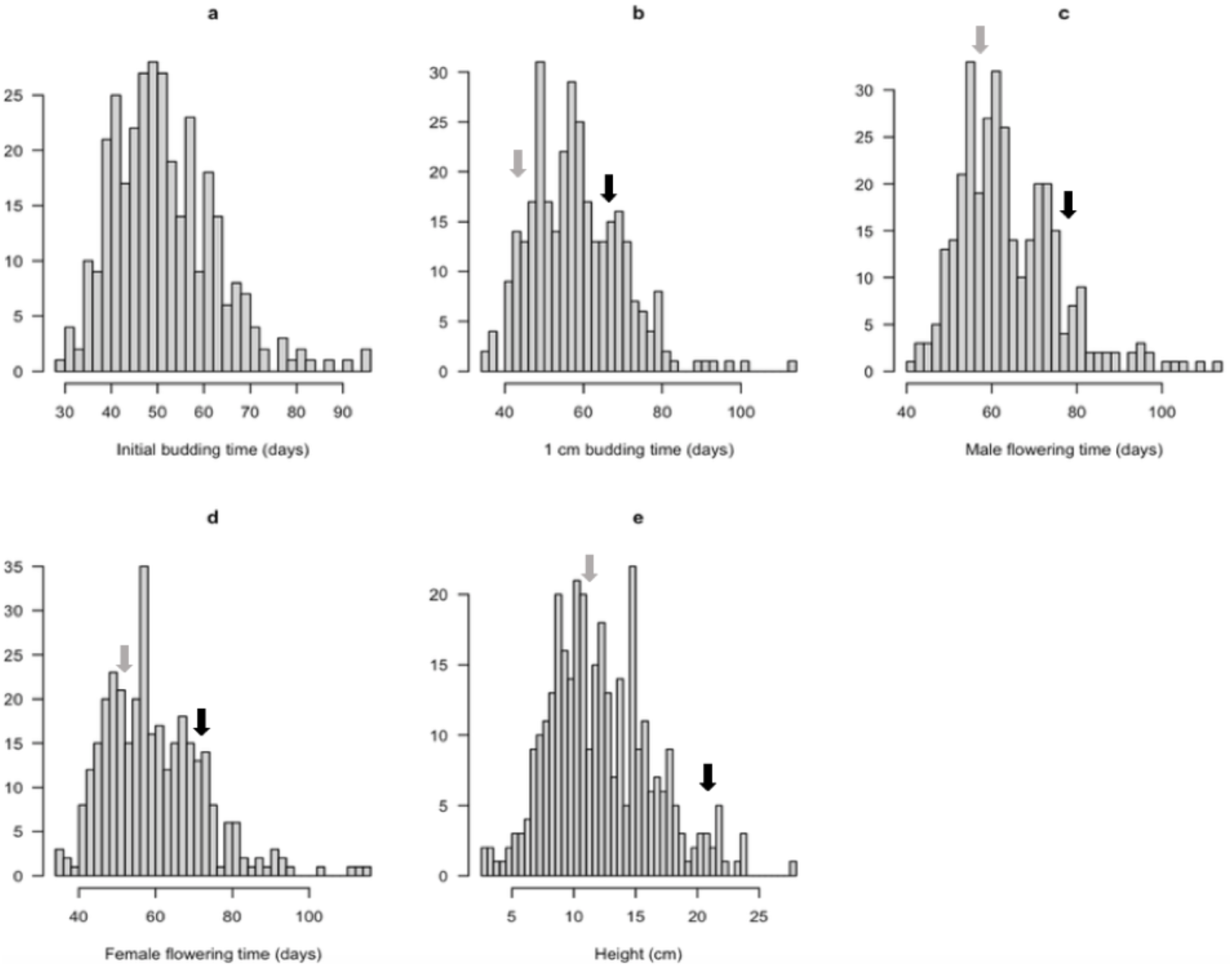
Histograms showing the frequency distributions of a) initial budding time b) 1cm budding time c) male flowering time d) female flowering time and e) height for the yellow mapping family of common ragweed (n=336). Grey arrows denote the phenotypic values of the early-flowering parent from the introduced European range. Black arrows denote the phenotypic values of the late-flowering parent from the native North American range. Initial budding time was not measured in the parental generation.

Marker orders within LGs were further checked using a crossover count method. This method involves counting the number of obligate crossovers for each possible order to find the order with the minimum number of crossovers (Broman & Sen, 2009). A large window size of seven markers was used first, followed by a smaller window of three, yet no orders were more likely than the original. To test for unusually tight double-crossovers, genotyping error LOD scores for all individuals at each marker were calculated (a LOD score higher than a specified cut-off indicates potential genotyping errors). There were no error LOD scores above a specified cutoff of four, three or two indicating no important errors of this kind. Finally, individual crossover counts were checked and a scatterplot of the observed number of crossovers revealed no major outliers. The final cross data used for QTL analysis consisted of phenotype data for 336 individuals, and their genotypes at 3,995 markers.

### QTL scans

Firstly, the function *calc.genoprob* was used to calculate QTL genotype probabilities conditional on the observed marker data, at a genotyping error rate of 0.01. Single-QTL scans with a normal model were subsequently performed for all five phenotypes using the function *scanone* and the extended Haley-Knott regression method (*method=ehk*). This method was chosen as it is an improved version of Haley–Knott regression (i.e., offers better approximation) and is more robust than standard interval mapping (Broman & Sen, 2009). For each scan, individuals with missing phenotype data were removed by the program (between 3-18 individuals per phenotype). We performed permutation tests (x1000) to obtain 0.1% and 5% genome-wide LOD significance thresholds, which were applied to each LOD distribution curve. The function *bayesint* was then used to derive Bayesian credible confidence intervals for the locations of identified QTLs at a coverage probability of 95%. To identify multiple pairs of linked or potentially interacting QTL, two dimensional two-QTL scans were carried out using the *scantwo* function. For each phenotype, permutation tests (x1000) were run to generate 5% genome-wide LOD significance thresholds for full, conditional-interactive, interactive, additive, and conditional-additive models. Using the results from *scanone* and *scantwo* and drawing from 1000 genotype simulation replicates, a multiple-QTL model was fit for each phenotype using the function *fitqtl*. Within this same function, estimates of QTL effect-sizes were obtained through analysis of variance (ANOVA). The integrated, high-density linkage map and QTLs were visualised using the R/LinkageMapView package (Ouellette, Reid, Blanchard, & Brouwer, 2017).

### Identifying candidate genes

To identify homologous flowering time genes within each QTL interval, we conducted BLAST searches of 29,849 annotated ragweed gene models (Battlay *et al*. in prep.) against *Arabidopsis* proteins with an *E*-value cut off of 1×10^-6^. The putative function of genes with a top hit in *Arabidopsis* were identified using gene ontology (GO) terms from The Arabidopsis Information Resource (TAIR) database (Berardini et al., 2015). Genes that had annotations relating to photoperiodism, floral development/regulation, vernalisation and circadian rhythm were flagged as potential candidates. Additionally, annotations were cross-referenced with 306 *A. thaliana* flowering time pathway genes (Bouché, Lobet, Tocquin, & Périlleux, 2016). 538 predicted *A. artemisiifolia* genes were matched to this dataset, representing 212 unique *A. thaliana* flowering time genes.

## Results

### Linkage map

The final integrated linkage map consisted of 4,493 markers spread over 18 linkage groups with a total length of 1,825.7 cM (Figure 2). LGs ranged in size from 55.1 cM to 139.4 cM with an average length of 101.4 cM. The average genetic distance between markers was 3.25 cM, with the largest gap being 13.34 cM in length (Figure 2).

**Figure 2.**
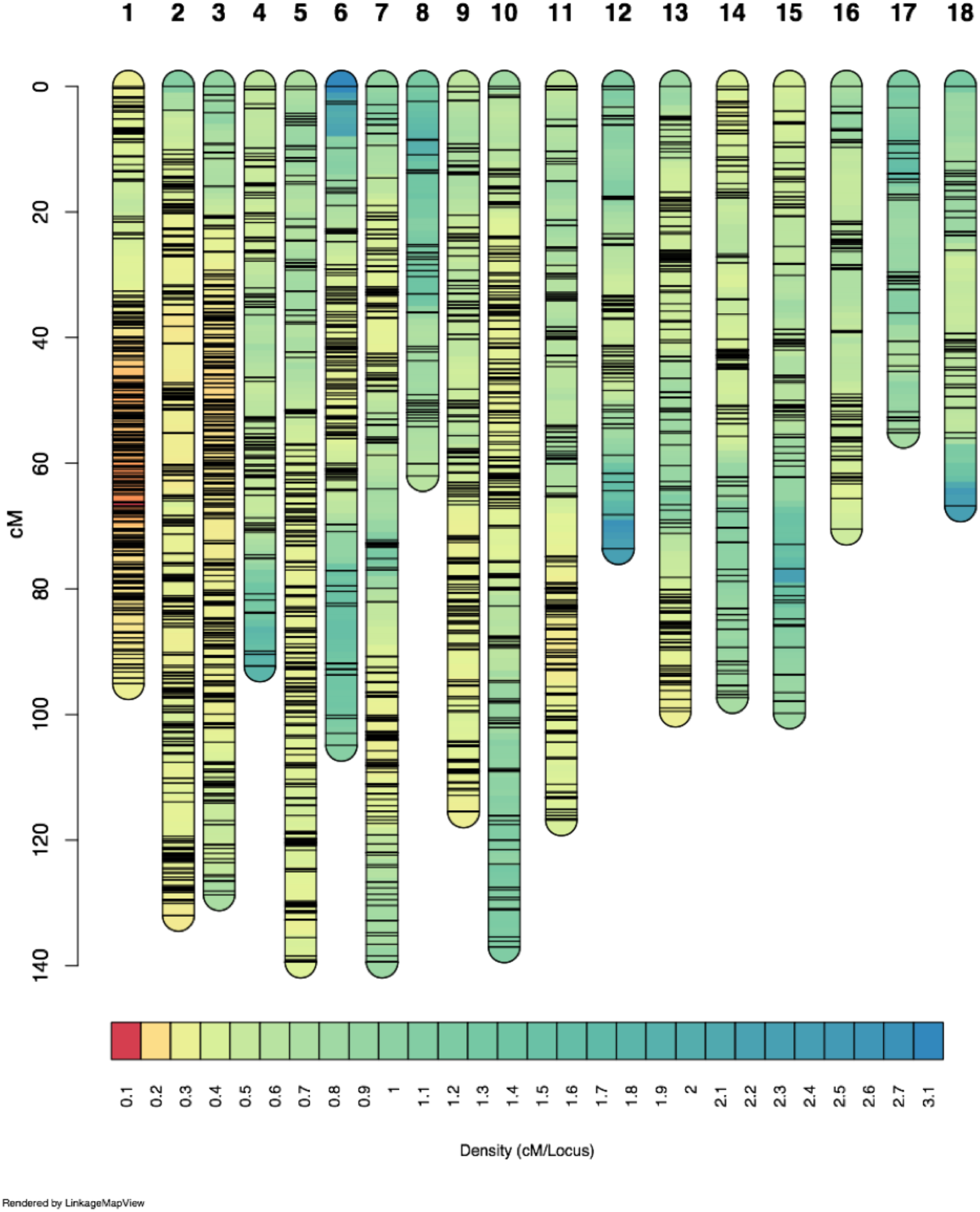
Density chart showing linkage group lengths and marker distributions of the integrated linkage map constructed from two mapping families of common ragweed. The 18 linkage groups correspond to the study system’s karyotype (2n = 36).

### QTL analysis

Single-QTL scans detected a significant QTL on LG2 (*QTL-2*) and LG12 (*QTL-12*) for all budding and flowering time traits (Figure 3; Table 1). A significant QTL for height was also found to co-localise with the flowering traits on LG2, with LOD scores maximised on the same marker but with a wider interval estimate of 82.2 cM compared to 13.2-23.0 cM (Table 1). Interval estimates for flowering and budding traits on *QTL-12* were relatively small, spanning 1.8-7.8cM (Table 1). Two-dimensional two-QTL scans did not identify any significant QTL-QTL interactions. Instead, the previously identified QTL on LG2 and LG12 were recognised as a significant pair of additive QTL. An additional QTL on LG6 (*QTL-6*) was found to have an additive interaction with *QTL-2* but only for female flowering time (Table 1). The interval estimate for this additional QTL spanned 38.1 cM (Table 1). *QTL-2* was a major QTL for budding and flowering traits, explaining 22.12-23.16% of observed phenotypic variance (Table 1). For height, *QTL-2* was defined as a minor QTL as it explained less than 10% of the phenotypic variation (Table 1). *QTL-12* was defined as a minor-effect QTL for budding and flowering traits, explaining 5.13-6.70% of phenotypic variance (Table 1). *QTL-6* was also of minor effect, explaining 2.73% of the variation in female flowering time (Table 1).

**Figure 3.**
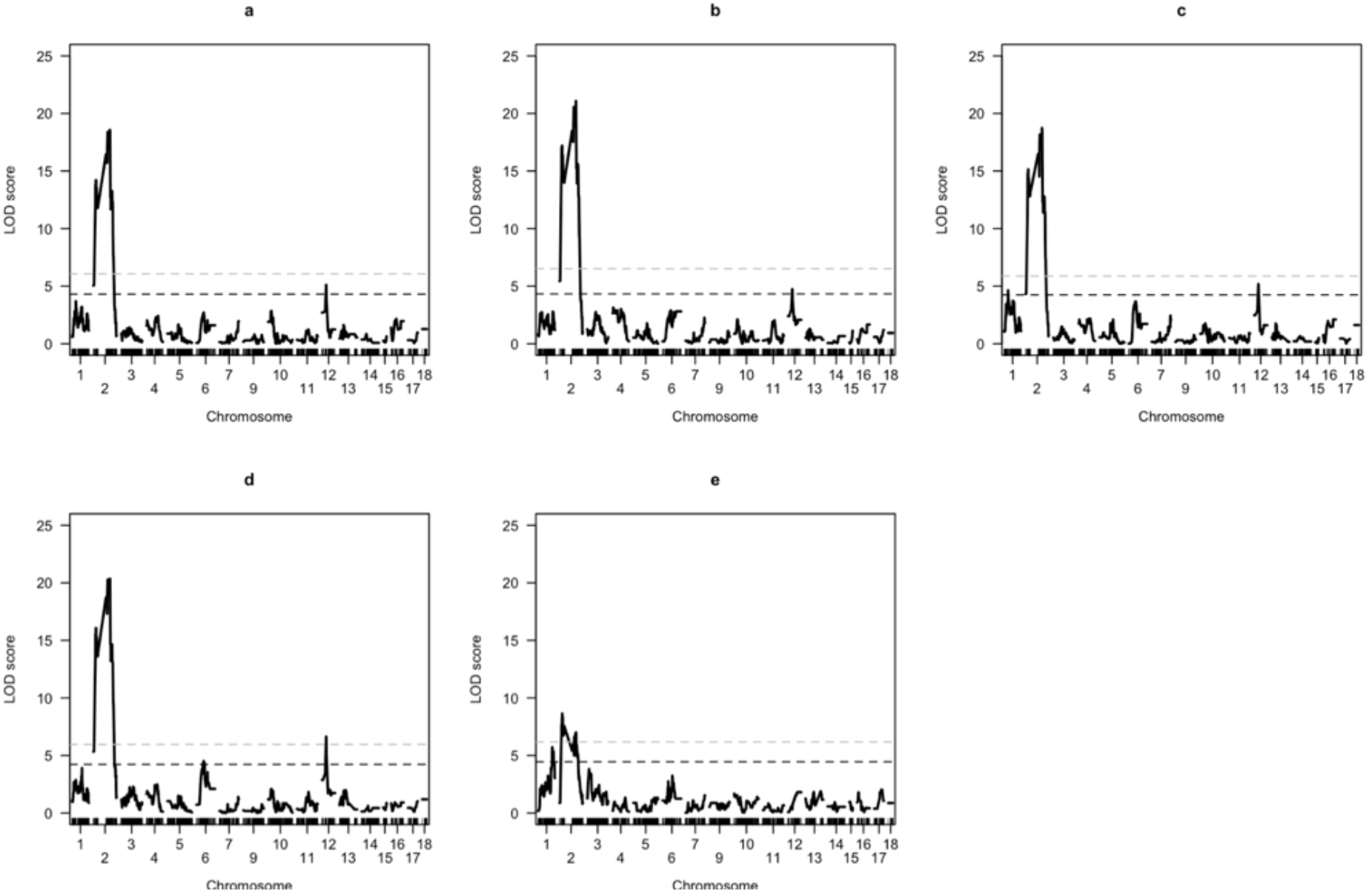
LOD distribution curves for a) initial budding time b) 1cm budding time c) male flowering time d) female flowering time and e) height, based on single-QTL scans of an experimental mapping population of common ragweed. Dotted lines represent genome-wide significance thresholds at 0.1% (light-grey dashes) and 5% (dark-grey dashes).

**Figure 4.**
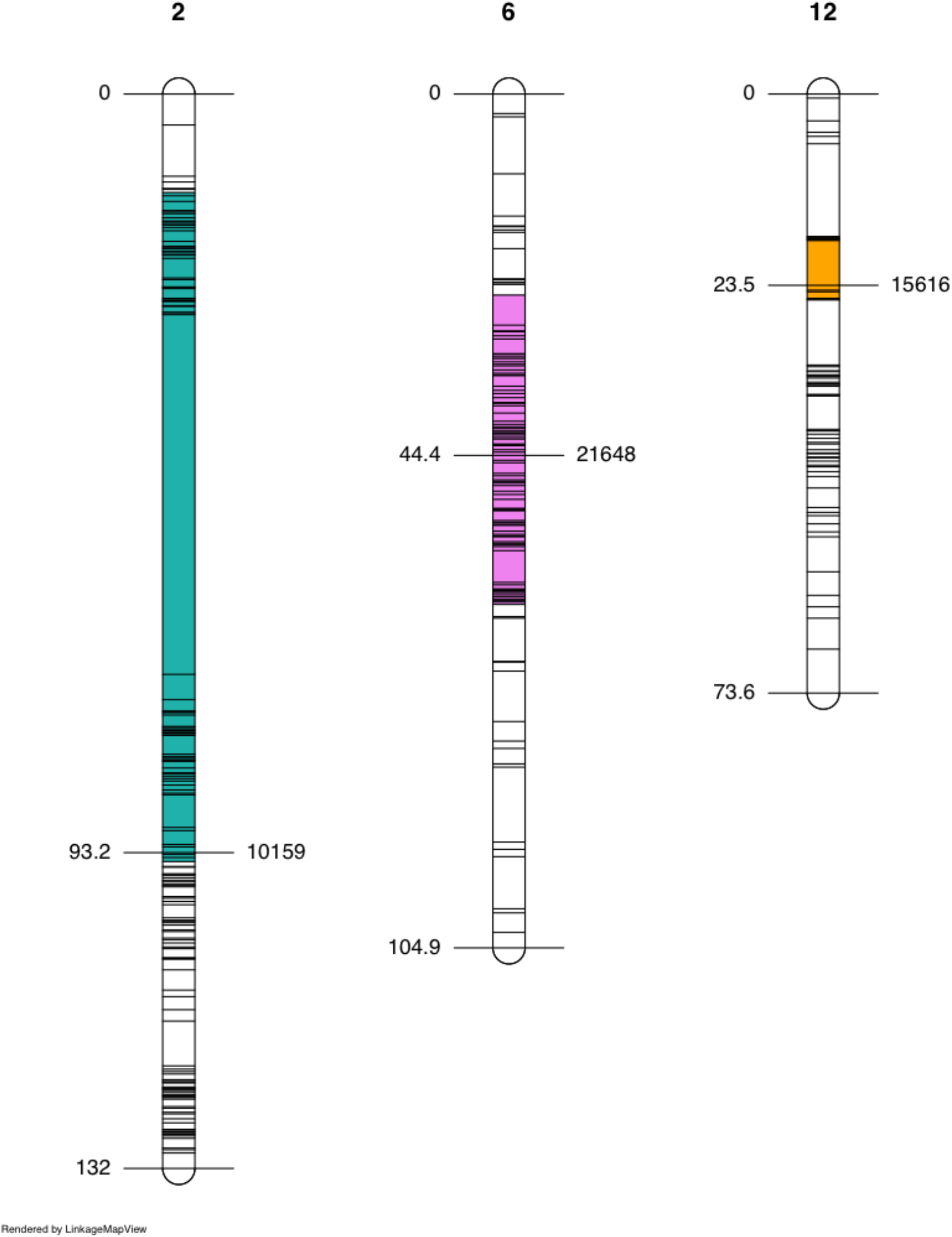
Linkage groups 2, 6 and 12 and their corresponding quantitative trait loci (QTL). Central markers where LOD scores were maximised are to the right and their genomic positions in centimorgans (cM) are to the left. Combined 95% Bayesian confidence intervals for all five traits are indicated by the colours sea-green (*QTL-2*), pink (*QTL-6*) and orange (*QTL-12*).

**Table 1.** Summary of QTL detected for flowering time and height in a common ragweed experimental mapping population. LG=linkage group. CI=confidence interval. % PVE=percentage of phenotypic variance explained by the QTL. Highly significant values (<0.001) are marked with ** and significant values (<0.05) are marked with *.

| QTL | LG | closest marker | position (cM) | 95% CI (cM) | trait | LOD | p-value | PVE (%) |
| --- | --- | --- | --- | --- | --- | --- | --- | --- |
| *QTL-2* | 2 | 10159 | 93.2 | 74.4-94.9 | initial budding time | 18.58 | <0.001** | 22.12 |
|  |  |  |  | 81.1-94.9 | 1 cm budding time | 19.81 | <0.001** | 22.12 |
|  |  |  |  | 81.1-94.3 | male flowering time | 18.74 | <0.001** | 22.1 |
|  |  |  |  | 71.3-94.3 | female flowering time | 20.35 | <0.001** | 23.16 |
|  |  |  |  | 12.1-94.3 | height (cm) | 6.687 | 0.001** | 6.7 |
| *QTL-12* | 12 | 15616 | 23.5 | 23.5-25.3 | initial budding time | 5.11 | 0.01* | 6.1 |
| *QTL-6* | 6 | 21648 | 44.4 | 17.5-25.3 | 1 cm budding time | 4.74 | 0.025* | 6.1 |
|  |  |  |  | 23.5-24.3 | male flowering time | 5.18 | 0.006* | 6.04 |
|  |  |  |  | 23.5-24.3 | female flowering time | 6.64 | 0.001** | 5.13 |
|  |  |  |  | 24.7-62.8 | female flowering time | 5.14 | 0.023* | 2.73 |

### Candidate flowering time genes

Within the *QTL-2* interval, nine genes with a top hit for *Arabidopsis* proteins had annotations relating to photoperiodism, circadian rhythm and flowering time regulation (Table 2), including two MADs box proteins closely related to FLOWERING LOCUS C (FLC). Only one gene had annotations relating to flowering time within the *QTL-12* interval (Table 2), and this was later identified as a homolog of FLOWERING LOCUS T (FT). Within the QTL interval on LG6, another nine genes were returned with annotations relating to photoperiodic sensing, circadian clock and flower development (Table 2).

**Table 2.**
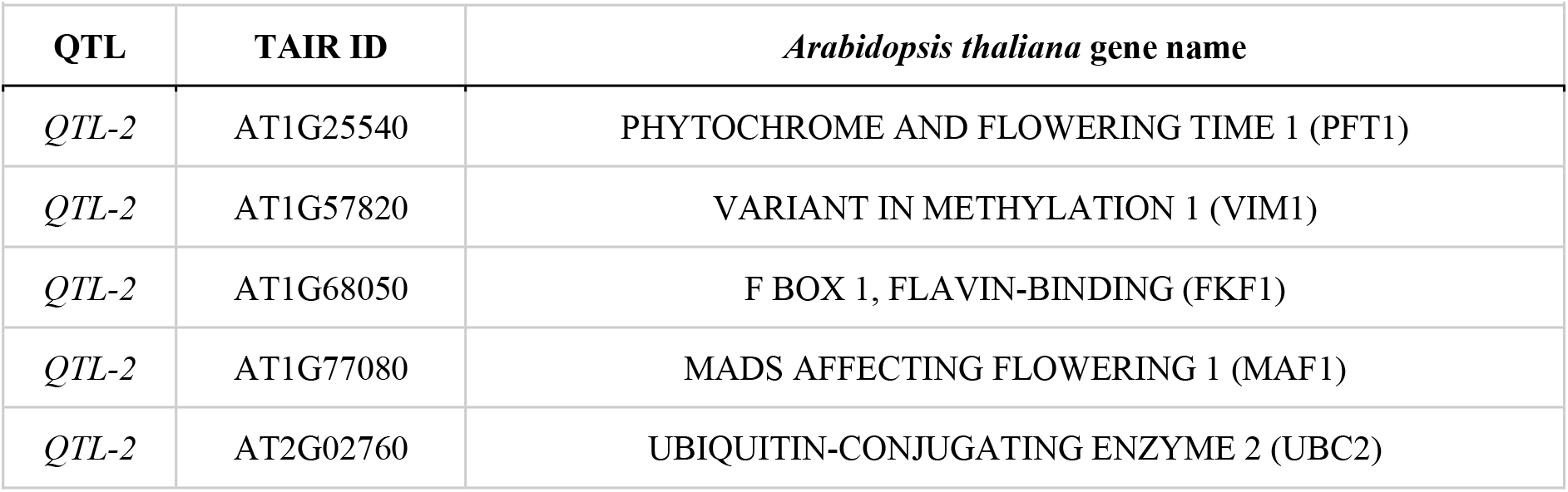
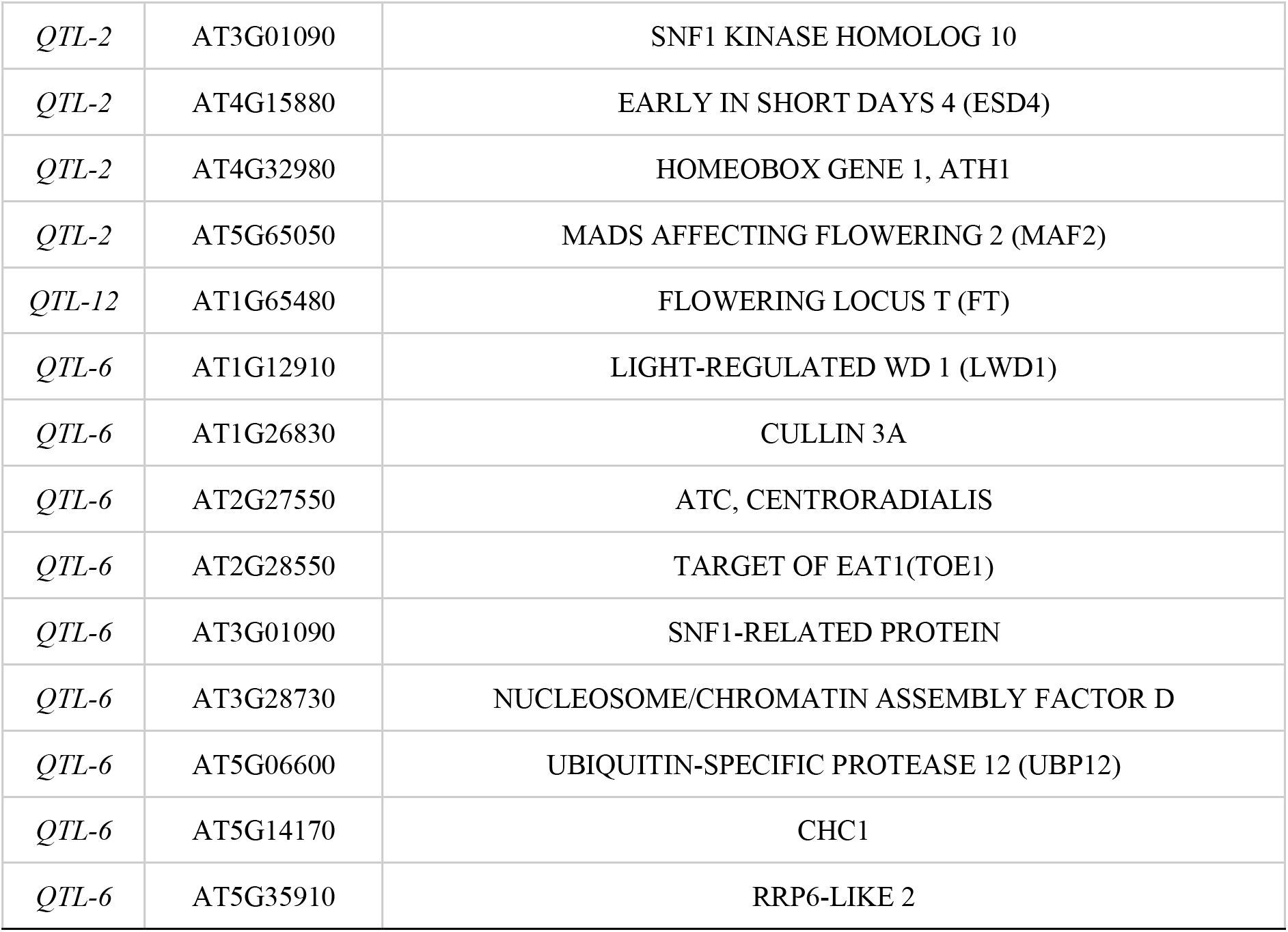
Flowering time candidates in each QTL-interval identified from BLAST searches between *A. thaliana* proteins and annotated common ragweed gene models.

### Recombination within QTL-associated haploblocks

*QTL-2* (Scaffold 27), which was associated with height and different measures of flowering time, colocalized with previously identified haploblocks (Battlay *et al*. in prep.). As haploblocks can be produced via suppressed recombination among specific haplotypes, as would be expected in the case of inversions, we examined recombination along this scaffold in separate genetic maps for each family and sex (Figure 5). Haploblock HB27a overlapped with the large interval for height. We found evidence for reduced recombination across the entirety of haploblock HB27a in three maps from the yellow and pink family (Figure 5 a-c). In the male map of the yellow family and in both F1 maps (Figure 5 d-f), recombination within the 27a haploblock was observed, suggesting that these parents were homozygous for the haploblock haplotypes while the others were heterozygous. Haploblock HB27b colocalized with the intervals for flowering time and height. In the pink and the yellow family, recombination was strongly reduced in the first half of HB27b (Figure 5 a-d), which overlapped with the flowering time QTL. The remainder of the haploblock had substantial recombination in the pink family and in the female map for the yellow family, but limited evidence of recombination in the male map for the yellow family (Figure 5 c), suggesting this parent may have been heterozygous for the haploblock. The F1 map showed recombination throughout the region suggesting both parents were homozygous for haploblock HB27b.

**Figure 5.**
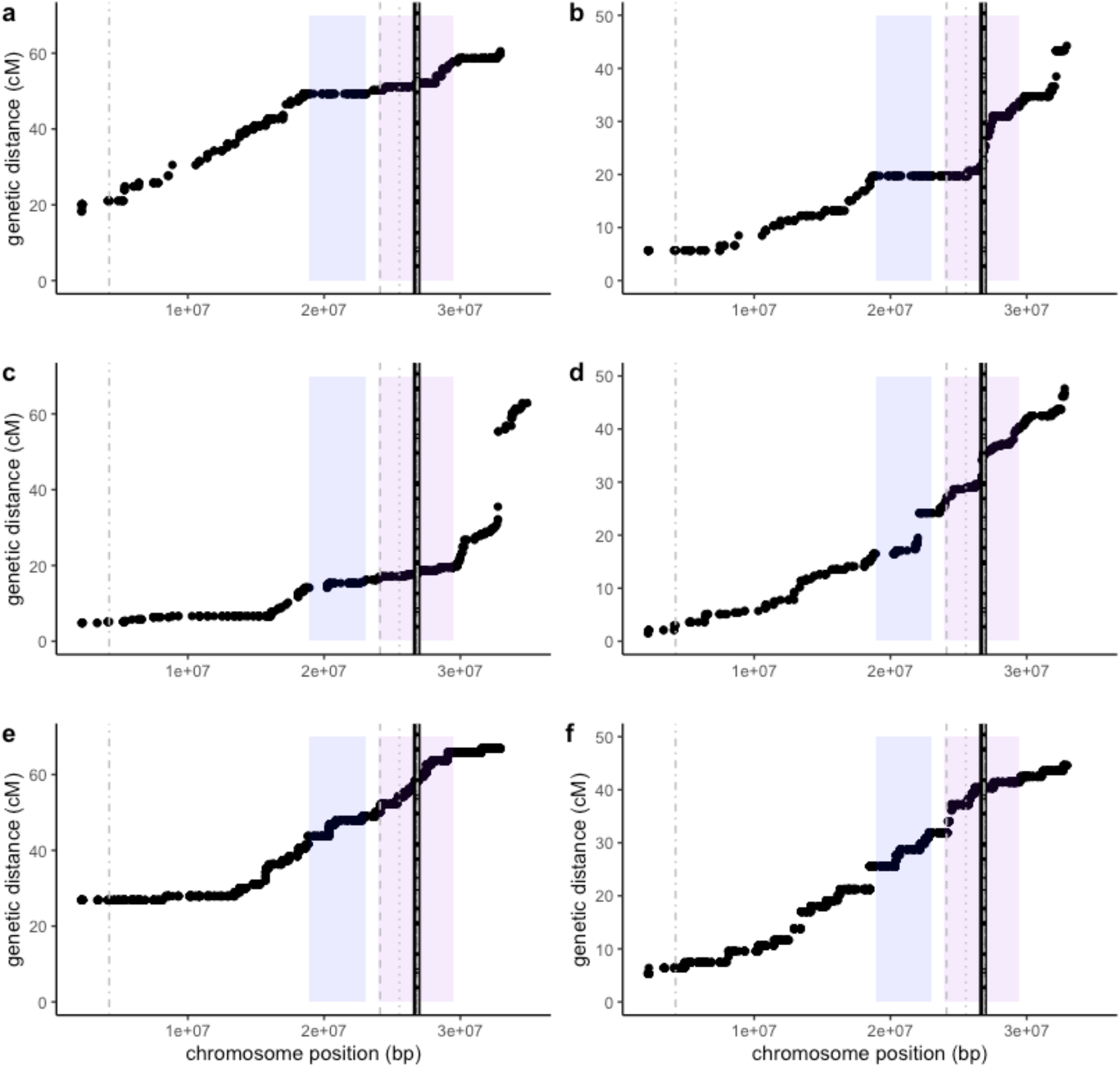
The genetic distance (cM) versus physical distance (bp) along a portion of Scaffold 27 containing *QTL-2*. Recombination distance was calculated for the pink (top), yellow (middle) and F1 families (bottom), and separately for each sex (male left; female right). The marker closest to *QTL-2* is represented by the black line. The 95% confidence intervals for male flowering time (dot), female flowering time (dash) and height (dash and dot) are shown in grey. The haploblocks are shaded (HB27a blue; HB27b purple).

## Discussion

Despite a growing interest in invasion biology from an evolutionary perspective, the genetic basis of rapid climate adaptation remains unresolved. To help address this knowledge gap, the present study attempted to dissect the genetic architecture underlying flowering time adaptation in the prolific invader, common ragweed. We identified one large-effect and two small-effect QTL underlying flowering time and height differentiation in this species. This is consistent with the expectation that when populations evolve under spatially divergent selection with migration, an oligogenic architecture is favoured. The co-localisation of flowering time and height QTL further supports the notion that gene pleiotropy is contributing to a genetic correlation (trade-off) between the two traits and the clinal variation in flowering time and height observed in common ragweed. Additionally, two putative inversions that overlap with *QTL-2* may be contributing to the accelerated local adaptation of this species observed in Europe. We also identified several candidate flowering time genes identified within the QTL that could be contributing to flowering time adaptation by controlling photoperiodic sensing and cold-repressed flowering. Together, these findings provide crucial insights into the genetic basis of climate adaptation during invasion, knowledge that is becoming increasingly invaluable as we strive to develop a more predictive understanding of species’ invasions in light of anthropogenic climate change.

### The genetic architecture of adaptation

Our discovery of one major and two minor QTL partly aligns with the theoretical prediction that adaptive evolution to local environments follows a stepwise approach to the optimum (Orr, 1998). In the early stages of an ‘adaptive walk’, when a population is far from its optimum (e.g., following colonisation of a new environment), both large and small-effect loci work in combination to produce large trait shifts which help to rapidly move the population closer to the phenotypic optimum (Dittmar, Oakley, Conner, Gould, & Schemske, 2016; Orr, 1998). However, as the population approaches the optimum, there is an increased likelihood that large-effect loci will reduce fitness by overshooting the optimum (Dittmar et al., 2016). At this point, small-effect loci that confer subtle trait shifts are required (Baxter, Johnston, & Jiggins, 2009; Orr, 1998). This leads to an exponential distribution of allele effect-sizes, with few loci of relatively large-effect and many of relatively small-effect (Dittmar et al., 2016; Orr, 1998). While this model may be applicable to some species (Bainbridge et al., 2020; McKay et al., 2008), we must take into account the effects of gene flow before we can confidently apply it to common ragweed. In contrast to this exponential distribution of allele effect sizes expected under adaptation to a single optimum without migration (Orr, 2005), divergent selection with migration is predicted to result in genetic architectures with fewer, larger, and more tightly linked alleles (i.e., oligogenic architecture) (Ferris, Barnett, Blackman, & Willis, 2017; Tigano & Friesen, 2016; Yeaman & Otto, 2011). Since large-effect loci tend to have larger selection coefficients compared to small-effect loci, they are more likely to be maintained by local selection despite the homogenising effects of gene flow (Yeaman & Otto, 2011). As such, prolonged bouts of adaptation in the face of ongoing gene flow could result in the gradual replacement of many small-effect alleles by fewer large-effect alleles, resulting in a genetic architecture that is skewed towards larger-effect loci (Yeaman & Otto, 2011; Yeaman & Whitlock, 2011). In some cases, these large-effect loci may often comprise several tightly linked genes each contributing small individual phenotypic effects (Yeaman & Whitlock, 2011). As an obligately outcrossing species reliant on wind pollination (Essl et al., 2015; Friedman & Barrett, 2008), common ragweed exhibits substantial gene flow across much of its native and introduced range (McGoey et al., 2020; van Boheemen et al., 2017). Hence, this high level of gene flow among ragweed’s divergently adapting populations may also serve to explain the oligogenic architecture observed in this QTL mapping population.

As a result of the interplay between divergent natural selection and migration, a genetic architecture that minimises recombination between locally adapted alleles is expected to evolve (Kirkpatrick & Barton, 2006; Lenormand & Otto, 2000; Yeaman & Whitlock, 2011). This is where the adaptation with gene flow theory dovetails neatly with the inversion literature (Ferris et al., 2017). When chromosomal inversions are heterozygous, they can suppress recombination and protect co-selected alleles from the hampering effects of gene flow (Wellenreuther, Rosenquist, Jaksons, & Larson, 2017). If the alleles captured inside the inversion afford the bearer greater fitness than alternative allelic combinations, the inversion can spread through a population and drive local adaptation (Kirkpatrick & Barton, 2006). Indirect evidence that inversions facilitate local adaptation stems from studies showing clines of inversion frequencies along environmental gradients (Day, Dawe, Dobson, & Hillier, 1983), and from other research linking inversions with adaptive functional variation in organisms such as fish (Kirubakaran et al., 2016) and flies (Ayala, Ullastres, & González, 2014). In plants, chromosomal inversions have frequently been associated with flowering time, which can contribute to both local adaptation and assortative mating (reviewed in (Huang & Rieseberg, 2020)). For example, (Lowry & Willis, 2010) demonstrated that a chromosomal inversion associated with flowering time in yellow monkeyflower (*Mimulus guttatus*) contributed to an adaptive annual-perennial life-history shift and reproductive isolation in this species. Evidently, chromosomal inversions are a potent local adaptation mechanism and can be maintained despite high gene flow.

In the present study, *QTL-2* overlaps with two large, divergent haplotype blocks or “haploblocks”, which indicate reduced recombination indicative of chromosomal inversions (Battlay *et al*. in prep.). *QTL-2* for height overlaps both haploblocks (HB27a and HB27b), while the *QTL-2* for flowering time-related traits have smaller intervals that only overlap the first half of HB27b (Figure 5). Our pink and yellow genetic maps reveal substantial reductions in recombination within HB27a (overlapping the height QTL) and in the first half of HB27b (overlapping the height and flowering time QTL). Recombination also appears to be reduced in the male map for the yellow family across all of HB27b (Figure 5c), which is the family used for the trait mapping. This suggests that the haploblock haplotypes are contributing to the large-effect QTL for flowering time which we identified in this study. The fact that some maps show repressed recombination while others don’t indicate these haploblocks are not located in regions with generally low recombination, but rather that reduced recombination is dependent on the genotypes used in the cross. This would occur, for example, if inversions had suppressed recombination when heterozygous, but not when homozygous. Although the evidence is largely indirect at this stage, these putative inversions could be capturing several tightly-linked locally adapted alleles, causing sections of *QTL-2* to act as single units with substantial effects on locally adapting traits including height and flowering time. In doing so, it could be creating a concentrated genetic architecture that is favoured under selection with gene flow and, in turn, may be contributing to the rapid local adaptation of common ragweed. Future studies confirming the mechanism underlying the suppressed recombination in these regions (e.g., inversions) and linking haplotypes to trait variation and fitness effects in divergent climates in the field will be important in addressing this hypothesis.

A potential caveat of this study is that the modest size of 336 F2s (from a single family) may not have allowed the detection of many small-effect QTL. This inherent bias towards the detection of large-effect QTL is common in QTL studies (Beavis, 2019; Remington, 2015; Rockman, 2012). Additionally, we only utilized one mapping family in our QTL analysis, as the others had little statistical power due to small sample size. Future studies could conduct QTL mapping on multiple and larger mapping families to improve resolution, and further interrogate the genetic basis of adaptation in a wider array of genotypes. Moreover, it is possible that additional QTL reside in the regions where markers had to be removed due to extreme segregation distortion, particularly on the linkage group where *QTL-2* was found. In fact, all markers within haploblock HB27a (in *QTL-2*) experienced segregation distortion in the yellow QTL mapping population. Segregation distortion is common in mapping studies, can be associated with chromosomal inversions (Fuller et al., 2020; X. Li, Wang, Wei, & Brummer, 2011), and is often caused by meiotic drive (Lyttle, 1993). Interestingly, HB27a is enriched for pectate lyase genes that are hypothesised to be involved in pollen tube growth (Chen, Marusciac, Tamas, Valenta, & Panaitescu, 2018), including the primary pollen allergen gene and five other paralogs of this gene (Battlay *et al*. in prep.). With that being said, the high-density linkage map and adequately-sized mapping population used for QTL analysis in this study did lead to the detection of some loci of small-effect and one locus of large-effect, thereby shedding light on the genetic architecture underlying rapid climate adaptation in common ragweed.

Based on overlapping credible intervals and similar LOD profiles, many of the QTL detected in this study were shared among traits. In particular, the co-localisation of a QTL for height and a QTL for flowering time is consistent with the expectation that these two traits will be at least partly genetically correlated (Colautti & Barrett, 2010; Kralemann et al., 2018). This may be due to linkage, or gene pleiotropy, where a single locus controls multiple phenotypes (Roff, 1995). In the latter case, a pleiotropic gene that determines the onset of flowering could also be controlling the plant’s level of vegetative growth (Zu & Schiestl, 2017). That is, if mutations that cause early flowering also result in reduced height due to a shift in the allocation of resources, the co-localisation of height and flowering time QTL is expected (Shen, Xiang, Xu, Ge, & Li, 2018). Pleiotropic effects of flowering time genes on growth have been frequently identified (Auge et al., 2019). This evidence of co-localised height and flowering time QTL also lends support to a recent study on common ragweed, which (by proxy measurement) discovered that widespread pleiotropy facilitates local adaptation in this species, particularly when populations were far from their selective optima (Hämälä, Gorton, Moeller, & Tiffin, 2020).

### Flowering time candidates

The transition to flowering is a well-orchestrated process that consists of hundreds of genes and transcription factors embedded in a complex network (Franks & Hoffmann, 2012; Putterill, Laurie, & Macknight, 2004; Wellmer & Riechmann, 2010). To ensure flowering occurs at the right time, plants must accurately perceive and process a range of environmental and internal cues (Putterill et al., 2004). In the model plant *Arabidopsis thaliana,* these signals are integrated through four main pathways (photoperiod, temperature, gibberellin and autonomous) and involve a number of genes that are presumably shared amongst most angiosperms (Michaels, 2009; Putterill et al., 2004; Wellmer & Riechmann, 2010). In this study, we discovered two MADs-box proteins closely related to FLOWERING LOCUS C (FLC) within the major-effect *QTL-2* interval. Known as MADS AFFECTING FLOWERING 1 (MAF1) and 2 (MAF2), these genes, like FLC, are floral repressors in *Arabidopsis* (Ratcliffe, Nadzan, Reuber, & Riechmann, 2001; Oliver J. Ratcliffe, Kumimoto, Wong, & Riechmann, 2003). Through the autonomous and vernalisation pathways, they are likely to act independently or downstream of FLC to delay flowering until after prolonged exposure to cold temperatures (Ratcliffe et al., 2001). In comparison to the late-flowering phenotype of winter-annual accessions of *A. thaliana*, weak FLC alleles have been linked to earlier flowering phenotypes in summer-annual accessions, enabling these populations to adapt and flower rapidly in the absence of vernalisation (Michaels, He, Scortecci, & Amasino, 2003). Although previously suspected of being a Brassicaceae-specific clade (De Bodt et al., 2003), recent studies have shown that FLC and MAF genes can act to regulate cold-repressed flowering in many other eudicots (P. A. Reeves et al., 2007; Tian et al., 2015). This study corroborates these findings by demonstrating that FLC-like genes can occur in plants from the Asteraceae family, including common ragweed.

In *A. thaliana*, FRIGIDA (FRI) maintains high levels of FLC mRNA which causes the downregulation of FLC targets (i.e. floral activators such as FLOWERING LOCUS T) to ultimately repress flowering (Gazzani, Gendall, Lister, & Dean, 2003). In conjunction with FRI, it is now speculated that there are other floral repressors that act to maintain FLC levels (Reeves, Murtas, Dash, & Coupland, 2002). Here, we detected a homolog of EARLY IN SHORT DAYS4 (ESD4) which has been shown to encode a novel regulator of FLC expression alongside another gene known as VERNALIZATION INDEPENDENCE4 (VIP4) (Reeves et al., 2002; H. Zhang & van Nocker, 2002). A mutant ESD4 allele has also been associated with an extreme early-flowering phenotype caused in part by a reduction in the expression level of FLC in *Arabidopsis* (Reeves et al., 2002).

As a major floral integrator, FLOWERING LOCUS T (FT) has a central position in the *A. thaliana* genetic network that regulates flowering time (Corbesier et al., 2007). It is expressed predominantly in the leaves and moves to the shoot apex where it interacts with the FLOWERING LOCUS D (FD) transcription factor to induce flowering (Corbesier et al., 2007; Jaeger & Wigge, 2007). Conversely, the closely-related TERMINAL FLOWER 1 (TFL1) gene functions antagonistically to repress flowering (Moraes, Dornelas, & Martinelli, 2019). This balance between FT/TFL1 is said to control photoperiodic flowering through competition with FD (Ahn et al., 2006), and is likely conserved between *A. thaliana* and other plant species (Higuchi et al., 2013). The discovery of an FT homolog within the *QTL-12* interval and a TFL1-like gene CENTRORADIALIS HOMOLOG (ATC) within the *QTL-6* interval suggests that the FT/TFL1 family of genes has conserved effects on floral induction in common ragweed. This is complementary to a previous study which revealed an association between differences in flowering time and variation in FT/TFL1-like gene expression levels in divergent populations of common ragweed (Kralemann et al., 2018). They found that earlier flowering in northern European populations was accompanied by a higher expression of an FT/TFL1 floral activator gene, enabling these populations to adapt to the shorter growing season (Kralemann et al., 2018). Notably, the divergent populations used in this expression analysis were the same ones used to generate the QTL mapping families in this study. This reaffirms the notion that the FT/TFL1 family of genes, and their effect on photoperiodic flowering, are highly conserved in common ragweed and are also likely to be major contributors to flowering time adaptation in this species.

We identified three homologs suspected of indirectly affecting FT transcription. Specifically, F-BOX1 (FKF1), PHYTOCHROME AND FLOWERING TIME 1 (PFT1) and TARGET OF EAT1 (TOE1). In the *A. thaliana* photoperiodic pathway, FKF1 has been shown to positively regulate the transcription factor CONSTANTS (CO), which is an FT promoter (Song, Smith, To, Millar, & Imaizumi, 2012). PFT1 has also been shown to activate CO and subsequent FT transcription (Iñigo, Alvarez, Strasser, Califano, & Cerdán, 2012). While initially identified as a nuclear protein responsible for the negative regulation of the phyB pathway (Cerdán & Chory, 2003), recent evidence has shown that the partial degradation of PFT1 facilitates FT transcription and promotes flowering in *A. thaliana* (Iñigo, Giraldez, Chory, & Cerdán, 2012). On the other hand, TOE1 and related proteins can bind near the activation domain of CO to ultimately inhibit CO and FT activity (Zhang, Wang, Zeng, Zhang, & Ma, 2015). Studies have also shown that TOE1 disturbs the FKF1–CO interaction by interfering with the LOV domain of FKF1 (Zhang et al., 2015). This likely results in the de-regulation of the CO protein in order to prevent premature flowering under suboptimal conditions (Zhang et al., 2015).

Overall, the homologs identified here indicate that variation of genes involved in the autonomous, vernalisation and photoperiodic pathways may allow common ragweed to time flowering in accordance with favourable climatic conditions. Their colocalization with the QTL identified here suggests at least some of these genes are involved in the flowering time differentiation of this species. In particular, future analysis of the function of natural variants within *QTL-2* will be important in assessing if single or multiple mutations in one or more flowering time genes are contributing to the large phenotypic effects we identified.

## Acknowledgements

We thank Carol Baskin and Andreas Lemke for providing seeds. This work was supported by grants from FORMAS to LA & RS (2016-00453) and by the ARC to KH (DP220102362 and DP180102531).

## Data accessibility

Sequence data will be available at the National Center for Biotechnology Information Sequence Read Archive under Bioproject XX. Raw phenotypic data will be available at XX. R scripts are available at https://github.com/khodgins/ragweed2021.

## References

Ågren, J., Oakley, C. G., Lundemo, S., & Schemske, D. W. (2017). Adaptive divergence in flowering time among natural populations of *Arabidopsis thaliana:* Estimates of selection and QTL mapping. Evolution; International Journal of Organic Evolution, 71(3), 550–564.

Ahn, J. H., Miller, D., Winter, V. J., Banfield, M. J., Lee, J. H., Yoo, S. Y.,… Weigel, D. (2006). A divergent external loop confers antagonistic activity on floral regulators FT and TFL1. The EMBO Journal, 25(3), 605–614.

Anderson, J. T., Willis, J. H., & Mitchell-Olds, T. (2011). Evolutionary genetics of plant adaptation. Trends in Genetics: TIG, 27(7), 258–266.

Auge, G. A., Penfield, S., & Donohue, K. (2019). Pleiotropy in developmental regulation by flowering-pathway genes: is it an evolutionary constraint? The New Phytologist, 224(1), 55–70.

Ayala, D., Ullastres, A., & González, J. (2014). Adaptation through chromosomal inversions in Anopheles. Frontiers in Genetics, 5, 129.

Bainbridge, H. E., Brien, M. N., Morochz, C., Salazar, P. A., Rastas, P., & Nadeau, N. J. (2020). Limited genetic parallels underlie convergent evolution of quantitative pattern variation in mimetic butterflies. Journal of Evolutionary Biology, 33(11), 1516–1529.

Bassett, I. J., & Crompton, C. W. (1975). The biology of Canadian weeds.: 11. *Ambrosia artemisiifolia* L. and *A. psilostachya* DC. Canadian Journal of Plant Science. Revue Canadienne de Phytotechnie, 55(2), 463–476.

Baxter, S. W., Johnston, S. E., & Jiggins, C. D. (2009). Butterfly speciation and the distribution of gene effect sizes fixed during adaptation. Heredity, 102(1), 57–65.

Beavis, W. D. (2019). 10 QTL Analyses: Power, Precision, and Accuracy. Molecular Dissection of Complex Traits. Retrieved from https://books.google.com/books?hl=en&lr=&id=IOavDwAAQBAJ&oi=fnd&pg=PT182&dq=QTL+Analyses:+Power,+Precision,+and+Accuracy.&ots=mEInOLOwey&sig=KJdfvXtIarsV0ZJjVOuK-aeL7PM

Berardini, T. Z., Reiser, L., Li, D., Mezheritsky, Y., Muller, R., Strait, E., & Huala, E. (2015). The arabidopsis information resource: Making and mining the “gold standard” annotated reference plant genome. Genesis, Vol. 53, pp. 474–485. doi: 10.1002/dvg.22877

Bieker, V. C., Battlay, P., Petersen, B., Sun, X., Wilson, J., Brealey, J. C.,… Martin, M. D. (2022). Uncovering the hologenomic basis of an extraordinary plant invasion. bioRxiv. doi: 10.1101/2022.02.03.478494

Bock, D. G., Caseys, C., Cousens, R. D., Hahn, M. A., Heredia, S. M., Hübner, S.,… Rieseberg, L. H. (2015). What we still don’t know about invasion genetics. Molecular Ecology, 24(9), 2277–2297.

Bouché, F., Lobet, G., Tocquin, P., & Périlleux, C. (2016). FLOR-ID: an interactive database of flowering-time gene networks in *Arabidopsis thaliana*. Nucleic Acids Research, 44(D1), D1167–D1171.

Broman, K. W., & Sen, S. (2009). A Guide to QTL Mapping with R/qtl. Springer, New York, NY.

Broman, K. W., Wu, H., Sen, S., & Churchill, G. A. (2003). R/qtl: QTL mapping in experimental crosses. Bioinformatics, 19(7), 889–890.

Bruelheide, H., & Heinemeyer, A. (2002). Climatic factors controlling the eastern and altitudinal distribution boundary of *Digitalis purpurea* L. in Germany. Flora - Morphology, Distribution, Functional Ecology of Plants, 197(6), 475–490.

Callaway, R. M., & Maron, J. L. (2006). What have exotic plant invasions taught us over the past 20 years? Trends in Ecology & Evolution, 21(7), 369–374.

Catchen, J., Hohenlohe, P. A., Bassham, S., Amores, A., & Cresko, W. A. (2013). Stacks: an analysis tool set for population genomics. Molecular Ecology, 22(11), 3124–3140.

Cerdán, P. D., & Chory, J. (2003). Regulation of flowering time by light quality. Nature, 423(6942), 881–885.

Chapman, D. S., Scalone, R., Štefanić, E., & Bullock, J. M. (2017). Mechanistic species distribution modeling reveals a niche shift during invasion. Ecology, 98(6), 1671–1680.

Chauvel, B., Dessaint, F., Cardinal-Legrand, C., & Bretagnolle, F. (2006). The historical spread of *Ambrosia artemisiifolia* L. in France from herbarium records. Journal of Biogeography, 33(4), 665–673.

Chen, K.-W., Marusciac, L., Tamas, P. T., Valenta, R., & Panaitescu, C. (2018). Ragweed pollen allergy: burden, characteristics, and management of an imported allergen source in Europe. International Archives of Allergy and Immunology, 176(3-4), 163–180.

Chun, Y. J., LE Corre, V., & Bretagnolle, F. (2011). Adaptive divergence for a fitness-related trait among invasive *Ambrosia artemisiifolia* populations in France. Molecular Ecology, 20(7), 1378– 1388.

Colautti, R. I., & Barrett, S. C. H. (2010). Natural selection and genetic constraints on flowering phenology in an invasive plant. International Journal of Plant Sciences, 171(9), 960–971.

Colautti, R. I., & Barrett, S. C. H. (2013). Rapid adaptation to climate facilitates range expansion of an invasive plant. Science, 342(6156), 364–366.

Colautti, R. I., & Lau, J. A. (2015). Contemporary evolution during invasion: evidence for differentiation, natural selection, and local adaptation. Molecular Ecology, 24(9), 1999–2017.

Collard, B. C. Y., Jahufer, M. Z. Z., Brouwer, J. B., & Pang, E. C. K. (2005). An introduction to markers, quantitative trait loci (QTL) mapping and marker-assisted selection for crop improvement: The basic concepts. Euphytica/Netherlands Journal of Plant Breeding, 142(1), 169–196.

Corbesier, L., Vincent, C., Jang, S., Fornara, F., Fan, Q., Searle, I.,… Coupland, G. (2007). FT protein movement contributes to long-distance signaling in floral induction of Arabidopsis. Science, 316(5827), 1030–1033.

Cunze, S., Leiblein, M. C., & Tackenberg, O. (2013). Range expansion of *Ambrosia artemisiifolia* in Europe is promoted by climate change. International Scholarly Research Notices, 2013. Retrieved from https://downloads.hindawi.com/archive/2013/610126.pdf

Danecek, P., Auton, A., Abecasis, G., Albers, C. A., Banks, E., DePristo, M. A.,… 1000 Genomes Project Analysis Group. (2011). The variant call format and VCFtools. Bioinformatics, 27(15), 2156–2158.

Day, T. H., Dawe, C., Dobson, T., & Hillier, P. C. (1983). A chromosomal inversion polymorphism in Scandinavian populations of the seaweed fly, Coelopa frigida. Hereditas, 99(1), 135–145.

De Bodt, S., Raes, J., Florquin, K., Rombauts, S., Rouzé, P., Theissen, G., & Van de Peer, Y. (2003). Genomewide structural annotation and evolutionary analysis of the type I MADS-box genes in plants. Journal of Molecular Evolution, 56(5), 573–586.

Dittmar, E. L., Oakley, C. G., Conner, J. K., Gould, B. A., & Schemske, D. W. (2016). Factors influencing the effect size distribution of adaptive substitutions. Proceedings. Biological Sciences /The Royal Society, 283(1828). doi: 10.1098/rspb.2015.3065

Dlugosch, K. M., Anderson, S. R., Braasch, J., Cang, F. A., & Gillette, H. D. (2015). The devil is in the details: genetic variation in introduced populations and its contributions to invasion. Molecular Ecology, 24(9), 2095–2111.

Essl, F., Biró, K., Brandes, D., Broennimann, O., Bullock, J. M., Chapman, D. S.,… Follak, S. (2015). Biological flora of the British isles: Ambrosia artemisiifolia. The Journal of Ecology, 103(4), 1069–1098.

Ferris, K. G., Barnett, L. L., Blackman, B. K., & Willis, J. H. (2017). The genetic architecture of local adaptation and reproductive isolation in sympatry within the *Mimulus guttatus* species complex. Molecular Ecology, 26(1), 208–224.

Fisher, R. A. (1919). XV.—The Correlation between Relatives on the Supposition of Mendelian Inheritance. Earth and Environmental Science Transactions of the Royal Society of Edinburgh, 52(2), 399–433.

Franks, S. J., & Hoffmann, A. A. (2012). Genetics of climate change adaptation. Annual Review of Genetics, 46, 185–208.

Franks, S. J., & Munshi-South, J. (2014). [Review of *Go forth, evolve and prosper: the genetic basis of adaptive evolution in an invasive species]*. Molecular ecology, 23(9), 2137–2140. Wiley Online Library.

Franks, S. J., Wheeler, G. S., & Goodnight, C. (2012). Genetic variation and evolution of secondary compounds in native and introduced populations of the invasive plant *Melaleuca quinquenervia*. Evolution; International Journal of Organic Evolution, 66(5), 1398–1412.

Fraser, D. J., Weir, L. K., Bernatchez, L., Hansen, M. M., & Taylor, E. B. (2011). Extent and scale of local adaptation in salmonid fishes: review and meta-analysis. Heredity, 106(3), 404–420.

Friedman, J., & Barrett, S. C. H. (2008). High outcrossing in the annual colonizing species *Ambrosia artemisiifolia* (Asteraceae). Annals of Botany, 101(9), 1303–1309.

Fuller, Z. L., Koury, S. A., Leonard, C. J., Young, R. E., Ikegami, K., Westlake, J.,… Phadnis, N. (2020). Extensive recombination suppression and epistatic selection causes chromosome-wide differentiation of a selfish sex chromosome in *Drosophila pseudoobscura*. Genetics, 216(1), 205–226.

Gazzani, S., Gendall, A. R., Lister, C., & Dean, C. (2003). Analysis of the Molecular Basis of Flowering Time Variation in Arabidopsis Accessions. Plant Physiology, 132(2), 1107–1114.

Gilchrist, G. W., & Lee, C. E. (2007). All stressed out and nowhere to go: does evolvability limit adaptation in invasive species? Genetica, 129(2), 127–132.

Gomulkiewicz, R., Holt, R. D., Barfield, M., & Nuismer, S. L. (2010). Genetics, adaptation, and invasion in harsh environments. Evolutionary Applications, 3(2), 97–108.

Griffith, T. M., & Watson, M. A. (2005). Stress avoidance in a common annual: reproductive timing is important for local adaptation and geographic distribution. Journal of Evolutionary Biology, 18(6), 1601–1612.

Grime, J. P. (1977). Evidence for the existence of three primary strategies in plants and its relevance to ecological and evolutionary theory. The American Naturalist, 111(982), 1169–1194.

Hämälä, T., Gorton, A. J., Moeller, D. A., & Tiffin, P. (2020). Pleiotropy facilitates local adaptation to distant optima in common ragweed (*Ambrosia artemisiifolia*). PLoS Genetics, 16(3), e1008707.

Helliwell, E. E., Faber-Hammond, J., Lopez, Z. C., Garoutte, A., von Wettberg, E., Friesen, M. L., & Porter, S. S. (2018). Rapid establishment of a flowering cline in *Medicago polymorpha* after invasion of North America. Molecular Ecology, 27(23), 4758–4774.

Hereford, J. (2009). A quantitative survey of local adaptation and fitness trade-offs. The American Naturalist, 173(5), 579–588.

Higuchi, Y., Narumi, T., Oda, A., Nakano, Y., Sumitomo, K., Fukai, S., & Hisamatsu, T. (2013). The gated induction system of a systemic floral inhibitor, antiflorigen, determines obligate short-day flowering in chrysanthemums. Proceedings of the National Academy of Sciences of the United States of America, 110(42), 17137–17142.

Hoban, S., Kelley, J. L., Lotterhos, K. E., Antolin, M. F., Bradburd, G., Lowry, D. B.,… Whitlock, M. C. (2016). Finding the genomic basis of local adaptation: Pitfalls, practical solutions, and future directions. The American Naturalist, 188(4), 379–397.

Hodgins, K. A., Bock, D. G., & Rieseberg, L. H. (2018). Trait evolution in invasive species. Annual Plant Reviews Online, 459–496.

Hodgins, K. A., & Rieseberg, L. (2011). Genetic differentiation in life-history traits of introduced and native common ragweed (*Ambrosia artemisiifolia*) populations. Journal of Evolutionary Biology, 24(12), 2731–2749.

Hodgins, K. A., & Yeaman, S. (2019). Mating system impacts the genetic architecture of adaptation to heterogeneous environments. The New Phytologist, 224(3), 1201–1214.

Huang, K., & Rieseberg, L. H. (2020). Frequency, origins, and evolutionary role of chromosomal inversions in plants. Frontiers in Plant Science, 11, 296.

Iñigo, S., Alvarez, M. J., Strasser, B., Califano, A., & Cerdán, P. D. (2012). PFT1, the MED25 subunit of the plant Mediator complex, promotes flowering through CONSTANS dependent and independent mechanisms in *Arabidopsis*. The Plant Journal: For Cell and Molecular Biology, 69(4), 601–612.

Iñigo, S., Giraldez, A. N., Chory, J., & Cerdán, P. D. (2012). Proteasome-mediated turnover of Arabidopsis MED25 is coupled to the activation of FLOWERING LOCUS T transcription. Plant Physiology, 160(3), 1662–1673.

Jaeger, K. E., & Wigge, P. A. (2007). FT protein acts as a long-range signal in Arabidopsis. Current Biology: CB, 17(12), 1050–1054.

Kawecki, T. J., & Ebert, D. (2004). Conceptual issues in local adaptation. Ecology Letters, 7(12), 1225–1241.

Keller, S. R., & Taylor, D. R. (2008). History, chance and adaptation during biological invasion: separating stochastic phenotypic evolution from response to selection. Ecology Letters, 11(8), 852–866.

Kirkpatrick, M., & Barton, N. (2006). Chromosome inversions, local adaptation and speciation. Genetics, 173(1), 419–434.

Kirubakaran, T. G., Grove, H., Kent, M. P., Sandve, S. R., Baranski, M., Nome, T.,… Andersen, Ø. (2016). Two adjacent inversions maintain genomic differentiation between migratory and stationary ecotypes of Atlantic cod. Molecular Ecology, 25(10), 2130–2143.

Kralemann, L. E. M., Scalone, R., Andersson, L., & Hennig, L. (2018). North European invasion by common ragweed is associated with early flowering and dominant changes in FT/TFL1 expression. Journal of Experimental Botany, 69(10), 2647–2658.

Laaidi, M., Laaidi, K., Besancenot, J.-P., & Thibaudon, M. (2003). Ragweed in France: an invasive plant and its allergenic pollen. Annals of Allergy, Asthma & Immunology: Official Publication of the American College of Allergy, Asthma, & Immunology, 91(2), 195–201.

Leiblein-Wild, M. C., & Tackenberg, O. (2014). Phenotypic variation of 38 European *Ambrosia artemisiifolia* populations measured in a common garden experiment. Biological Invasions, 16(9), 2003–2015.

Leimu, R., & Fischer, M. (2008). A meta-analysis of local adaptation in plants. PloS One, 3(12), e4010.

Lenormand, T., & Otto, S. P. (2000). The evolution of recombination in a heterogeneous environment. Genetics, 156(1), 423–438.

Li, H., & Durbin, R. (2010). Fast and accurate long-read alignment with Burrows–Wheeler transform. Bioinformatics, 26(5), 589–595.

Li, H., Handsaker, B., Wysoker, A., Fennell, T., Ruan, J., Homer, N.,… 1000 Genome Project Data Processing Subgroup. (2009). The Sequence Alignment/Map format and SAMtools. Bioinformatics, 25(16), 2078–2079.

Li, H., Hearne, S., Bänziger, M., Li, Z., & Wang, J. (2010). Statistical properties of QTL linkage mapping in biparental genetic populations. Heredity, 105(3), 257–267.

Li, X.-M., She, D.-Y., Zhang, D.-Y., & Liao, W.-J. (2015). Life history trait differentiation and local adaptation in invasive populations of *Ambrosia artemisiifolia* in China. Oecologia, 177(3), 669–677.

Li, X., Wang, X., Wei, Y., & Brummer, E. C. (2011). Prevalence of segregation distortion in diploid alfalfa and its implications for genetics and breeding applications. TAG. Theoretical and Applied Genetics. Theoretische Und Angewandte Genetik, 123(4), 667–679.

Louthan, A. M., & Kay, K. M. (2011). Comparing the adaptive landscape across trait types: larger QTL effect size in traits under biotic selection. BMC Evolutionary Biology, 11, 60.

Lowry, D. B., & Willis, J. H. (2010). A widespread chromosomal inversion polymorphism contributes to a major life-history transition, local adaptation, and reproductive isolation. PLoS Biology, 8(9). doi: 10.1371/journal.pbio.1000500

Lyttle, T. W. (1993). Cheaters sometimes prosper: distortion of mendelian segregation by meiotic drive. Trends in Genetics: TIG, 9(6), 205–210.

MacPherson, A., & Nuismer, S. L. (2017). The probability of parallel genetic evolution from standing genetic variation. Journal of Evolutionary Biology, 30(2), 326–337.

McGoey, B. V., Hodgins, K. A., & Stinchcombe, J. R. (2020). Parallel flowering time clines in native and introduced ragweed populations are likely due to adaptation. Ecology and Evolution, 10(11), 4595–4608.

McGoey, B. V., & Stinchcombe, J. R. (2021). Introduced populations of ragweed show as much evolutionary potential as native populations. Evolutionary Applications, 14(5), 1436–1449.

McKay, J. K., Richards, J. H., Nemali, K. S., Sen, S., Mitchell-Olds, T., Boles, S.,… Juenger, T. E. (2008). Genetics of drought adaptation in Arabidopsis thaliana II. QTL analysis of a new mapping population, KAS-1 x TSU-1. Evolution; International Journal of Organic Evolution, 62(12), 3014–3026.

Michaels, S. D. (2009). Flowering time regulation produces much fruit. Current Opinion in Plant Biology, 12(1), 75–80.

Michaels, S. D., He, Y., Scortecci, K. C., & Amasino, R. M. (2003). Attenuation of FLOWERING LOCUS C activity as a mechanism for the evolution of summer-annual flowering behavior in Arabidopsis. Proceedings of the National Academy of Sciences of the United States of America, 100(17), 10102–10107.

Moraes, T. S., Dornelas, M. C., & Martinelli, A. P. (2019). FT/TFL1: Calibrating plant architecture. Frontiers in Plant Science, 10, 97.

Oduor, A. M. O., Leimu, R., & van Kleunen, M. (2016). Invasive plant species are locally adapted just as frequently and at least as strongly as native plant species. The Journal of Ecology, 104(4),957–968.

Orr, H. A. (1998). The population genetics of adaptation: The distribution of factors fixed during adaptive evolution. Evolution; International Journal of Organic Evolution, 52(4), 935–949.

Orr, H. A. (2005). The genetic theory of adaptation: a brief history. Nature Reviews. Genetics, 6(2), 119–127.

Ouellette, L. A., Reid, R. W., Blanchard, S. G., & Brouwer, C. R. (2017). LinkageMapView— rendering high-resolution linkage and QTL maps. Bioinformatics, 34(2), 306–307.

Putterill, J., Laurie, R., & Macknight, R. (2004). It’s time to flower: the genetic control of flowering time. BioEssays: News and Reviews in Molecular, Cellular and Developmental Biology, 26(4), 363–373.

Rastas, P. (2017). Lep-MAP3: robust linkage mapping even for low-coverage whole genome sequencing data. Bioinformatics, 33(23), 3726–3732.

Ratcliffe, O. J., Kumimoto, R. W., Wong, B. J., & Riechmann, J. L. (2003). Analysis of the *Arabidopsis* MADS AFFECTING FLOWERING gene family: MAF2 prevents vernalization by short periods of cold. The Plant Cell, 15(5), 1159–1169.

Ratcliffe, O. J., Nadzan, G. C., Reuber, T. L., & Riechmann, J. L. (2001). Regulation of flowering in *Arabidopsis* by an FLC homologue. Plant Physiology, 126(1), 122–132.

Reeves, P. A., He, Y., Schmitz, R. J., Amasino, R. M., Panella, L. W., & Richards, C. M. (2007). Evolutionary conservation of the FLOWERING LOCUS C-mediated vernalization response: evidence from the sugar beet (Beta vulgaris). Genetics, 176(1), 295–307.

Reeves, P. H., Murtas, G., Dash, S., & Coupland, G. (2002). early in short days 4, a mutation in Arabidopsis that causes early flowering and reduces the mRNA abundance of the floral repressor FLC. Development, 129(23), 5349–5361.

Remington, D. L. (2015). Alleles versus mutations: Understanding the evolution of genetic architecture requires a molecular perspective on allelic origins. Evolution; International Journal of Organic Evolution, 69(12), 3025–3038.

Rockman, M. V. (2012). The QTN program and the alleles that matter for evolution: all that’s gold does not glitter. Evolution; International Journal of Organic Evolution, 66(1), 1–17.

Roff, D. A. (1995). The estimation of genetic correlations from phenotypic correlations: a test of Cheverud’s conjecture. Heredity, 74(5), 481–490.

Sakai, A. K., Allendorf, F. W., Holt, J. S., Lodge, D. M., Molofsky, J., With, K. A.,… Weller, S. G. (2001). The Population Biology of Invasive Species. Annual Review of Ecology and Systematics, 32(1), 305–332.

Santangelo, J. S., Johnson, M. T. J., & Ness, R. W. (2018). Modern spandrels: the roles of genetic drift, gene flow and natural selection in the evolution of parallel clines. Proceedings. Biological Sciences / The Royal Society, 285(1878). doi: 10.1098/rspb.2018.0230

Savolainen, O., Lascoux, M., & Merilä, J. (2013). Ecological genomics of local adaptation. Nature Reviews. Genetics, 14(11), 807–820.

Sax, D. F., & Brown, J. H. (2000). The paradox of invasion. Global Ecology and Biogeography: A Journal of Macroecology, 9(5), 363–371.

Scalone, R., Lemke, A., Štefanić, E., Kolseth, A.-K., Rašić, S., & Andersson, L. (2016). Phenological Variation in *Ambrosia artemisiifolia* L. facilitates near future establishment at northern latitudes. PloS One, 11(11), e0166510.

Shen, Y., Xiang, Y., Xu, E., Ge, X., & Li, Z. (2018). Major co-localized QTL for plant height, branch initiation height, stem diameter, and flowering time in an alien introgression derived *Brassica napus* DH population. Frontiers in Plant Science, 9, 390.

Sherrard, M. E., & Maherali, H. (2012). Local adaptation across a fertility gradient is influenced by soil biota in the invasive grass, *Bromus inermis*. Evolutionary Ecology, 26(3), 529–544.

Smith, M., Cecchi, L., Skjøth, C. A., Karrer, G., & Šikoparija, B. (2013). Common ragweed: a threat to environmental health in Europe. Environment International, 61, 115–126.

Song, Y. H., Smith, R. W., To, B. J., Millar, A. J., & Imaizumi, T. (2012). FKF1 conveys timing information for CONSTANS stabilization in photoperiodic flowering. Science, 336(6084), 1045– 1049.

Stinson, K. A., Wheeler, J. A., Record, S., & Jennings, J. L. (2018). Regional variation in timing, duration, and production of flowers by allergenic ragweed. Plant Ecology, 219(9), 1081–1092.

Sultan, S. E., Horgan-Kobelski, T., Nichols, L. M., Riggs, C. E., & Waples, R. K. (2013). A resurrection study reveals rapid adaptive evolution within populations of an invasive plant. Evolutionary Applications, 6(2), 266–278.

Taramarcaz, P., Lambelet, B., Clot, B., Keimer, C., & Hauser, C. (2005). Ragweed (*Ambrosia*)progression and its health risks: will Switzerland resist this invasion? Swiss Medical Weekly, 135(37-38), 538–548.

Tian, Y., Dong, Q., Ji, Z., Chi, F., Cong, P., & Zhou, Z. (2015). Genome-wide identification and analysis of the MADS-box gene family in apple. Gene, 555(2), 277–290.

Tigano, A., & Friesen, V. L. (2016). Genomics of local adaptation with gene flow. Molecular Ecology, 25(10), 2144–2164.

Todesco, M., Owens, G. L., Bercovich, N., Légaré, J.-S., Soudi, S., Burge, D. O.,… Rieseberg, L. H. (2020). Massive haplotypes underlie ecotypic differentiation in sunflowers. Nature, 584(7822), 602–607.

Tokarska-Guzik, B., Bzdęga, K., Koszela, K., Żabińska, I., Krzuś, B., Sajan, M., & Sendek, A. (2011). Allergenic invasive plant *Ambrosia artemisiifoli*a L. in Poland: threat and selected aspects of biology. Biodiversity Research and Conservation, 21(2011), 39–48.

van Boheemen, L. A., Atwater, D. Z., & Hodgins, K. A. (2019). Rapid and repeated local adaptation to climate in an invasive plant. The New Phytologist, 222(1), 614–627.

van Boheemen, L. A., & Hodgins, K. A. (2020). Rapid repeatable phenotypic and genomic adaptation following multiple introductions. Molecular Ecology, 29(21), 4102–4117.

van Boheemen, L. A., Lombaert, E., Nurkowski, K. A., Gauffre, B., Rieseberg, L. H., & Hodgins, K. A. (2017). Multiple introductions, admixture and bridgehead invasion characterize the introduction history of *Ambrosia artemisiifolia* in Europe and Australia. Molecular Ecology, 26(20), 5421–5434.

Weaver, S. E. (2001). Impact of lamb’s-quarters, common ragweed and green foxtail on yield of corn and soybean in Ontario. Canadian Journal of Plant Science. Revue Canadienne de Phytotechnie, 81(4), 821–828.

Wellenreuther, M., Rosenquist, H., Jaksons, P., & Larson, K. W. (2017). Local adaptation along an environmental cline in a species with an inversion polymorphism. Journal of Evolutionary Biology, 30(6), 1068–1077.

Wellmer, F., & Riechmann, J. L. (2010). Gene networks controlling the initiation of flower development. Trends in Genetics: TIG, 26(12), 519–527.

Westley, P. A. H. (2011). What invasive species reveal about the rate and form of contemporary phenotypic change in nature. The American Naturalist, 177(4), 496–509.

Yan, W., Wang, B., Chan, E., & Mitchell-Olds, T. (2021). Genetic architecture and adaptation of flowering time among environments. The New Phytologist, 230(3), 1214–1227.

Yeaman, S., & Otto, S. P. (2011). Establishment and maintenance of adaptive genetic divergence under migration, selection, and drift. Evolution; International Journal of Organic Evolution, 65(7), 2123–2129.

Yeaman, S., & Whitlock, M. C. (2011). The genetic architecture of adaptation under migration-selection balance. Evolution; International Journal of Organic Evolution, 65(7), 1897–1911.

Zhang, B., Wang, L., Zeng, L., Zhang, C., & Ma, H. (2015). Arabidopsis TOE proteins convey a photoperiodic signal to antagonize CONSTANS and regulate flowering time. Genes & Development, 29(9), 975–987.

Zhang, H., & van Nocker, S. (2002). The VERNALIZATION INDEPENDENCE 4 gene encodes a novel regulator of FLOWERING LOCUS C. The Plant Journal: For Cell and Molecular Biology, 31(5), 663–673.

Ziska, L., Knowlton, K., Rogers, C., Dalan, D., Tierney, N., Elder, M. A.,… Frenz, D. (2011). Recent warming by latitude associated with increased length of ragweed pollen season in central North America. Proceedings of the National Academy of Sciences of the United States of America, 108(10), 4248–4251.

Zu, P., & Schiestl, F. P. (2017). The effects of becoming taller: direct and pleiotropic effects of artificial selection on plant height in *Brassica rapa*. The Plant Journal: For Cell and Molecular Biology, 89(5), 1009–1019.

